# Gibbs Free Energy, a thermodynamic measure of protein-protein interactions, correlates with neurologic disability

**DOI:** 10.1101/527820

**Authors:** Michael Keegan, Hava T. Siegelmann, Edward A. Rietman, Giannoula Lakka Klement

## Abstract

**Background:** Modern network science has been used to reveal new and often fundamental aspects of brain network organization in physiological as well as pathological conditions. As a consequence, these discoveries, which relate to network hierarchy, hubs and network interactions, begun to change the paradigms of neurodegenerative disorders. We therefore explored the use of thermodynamics for protein-protein network interactions in Alzheimer disease (AD), Parkinson disease (PD), multiple sclerosis (MS), traumatic brain injury and epilepsy.

**Methods:** To assess the validity of using network interactions in neurological disease, we investigated the relationship between network thermodynamics and molecular systems biology for these neurological disorders. In order to uncover whether there was a correlation between network organization and biological outcomes, we used publicly available RNA transcription data from individual patients with these neurological conditions, and correlated these molecular profiles with their respective individual disability scores.

**Results:** We found a linear correlation (Pearson correlation of −0.828) between disease disability (a clinically validated measurement of a person’s functional status), and Gibbs free energy (a thermodynamic measure of protein-protein interactions). In other words, we found an inverse relationship between disease entropy and thermodynamic energy.

**Interpretation:** Because a larger degree of disability correlated with a larger negative drop in Gibbs free energy in a linear, disability-dependent fashion, it could be presumed that the progression of neuropathology such as is seen in Alzheimer Disease, could potentially be prevented by therapeutically correcting the changes Gibbs free energy.

## Background

The treatment and management of neurological dysfunction/ neurodegeneration is an area of a great medical need. The World Health Organization (WHO) estimates that neurological disorders contribute 10.9% and 8.7% of global disease burden in high- and medium-income countries,[1] and as the average age of the populations in developed countries increases in the coming decades, it is expected that the disease burden will continue to increase. Yet, the present treatment of neurological diseases constitutes by and large of management of disease symptoms because the etiology remains unclear. In Alzheimer disease for example, the earlier etiological hypothesis that amyloid deposits are caused by environmental stimuli is being supplanted by growing body of evidence that it is the genomic dysregulations of cellular and molecular pathways that cause accumulation of amyloid and tau proteins.[2] This may not only explain why targeting amyloid and/or tau proteins has been unsuccessful so far, it also unlocks the possibility of using targeted therapies. However, to realize the full potential of targeting these newly identified genomic patterns in Alzheimer Disease[3, 4] and in order to pursue these novel, molecularly-guided therapies, an improved understanding about the complexity of systems biology of neurological diseases is needed.

Many pathologic conditions such as for example neurodegenerative disease, chronic inflammatory disorders, or cancer; are associated not only with mutational activation of genes, but also with re-activation of developmentally silenced pathways. Because the processes of tissue invasion, proliferation, inflammation and angiogenesis are common not only to oncogenic induction, but also to embryonal development and normal host response to tissue injury - a simple DNA mutation analysis would not provide sufficient information. The activation of pathways associated with inflammation and tissue injury is therefore best studied using mRNA expression. However, until very recently, it has been very difficult to directly correlate the levels of gene expression with the cellular effect of specific proteins. In fact high gene expression did not always imply increased protein function or activation of a biological process. The intensity of a biological effect is dependent on the interaction of the affected (overexpressed) gene with its neighbors, and the quorum effect of the innumerable feedback loops on the global protein-protein interaction network.

The realization that the perturbations of individual genes can be measured by its effect on the global response of a network has led many scientists to finding alternative approaches for genomic interpretations. The one presented in this manuscript is based on using not only level of expression of a gene, but also its topology as a measure of its connectivity. This novel approach has been enabled by the vast amounts of well-curated information accumulated in publicly available protein-protein interaction networks (PPINs) over the last 4 decades, and has emerged quite recently.[5–8]

The underlying premise of our analysis is that biological systems are complex chemical networks. A cell, for example, consists of a large molecular network made of DNA, RNA, proteins, peptides, small molecules, and lipids. Each of these molecules is associated with potential energy contributing to an even larger energetic network. The energetic state exists in equilibrium, and any perturbance sets off a cascade of events striving to bring the overall network back to the same entropy. The prime force pushing the reaction back is Gibbs free energy, an expression of the thermodynamic energy reflecting the chemical potential between interacting proteins. This Gibbs Free Energy is used as a measure of the changes occurring within a disease-related protein-protein interaction network.

In his earlier work, Rietman *et al*[9] described an inverse correlation between Gibbs free energy and percent 5-yr survival for ten different types of cancers. The study showed that poor prognosis cancers such as glioblastoma multiforme have low thermodynamic entropy, less negative Gibbs free energy, and very low 5-year survival. This was consistent with the clinical status of the disease, which has average survival post diagnosis of 6 months, and a 5-yr survival of 2%. Similarly, the finding that in breast carcinoma, which had higher thermodynamic entropy, a more negative Gibbs free energy, and a much higher 5-year survival was also congruent with the clinical observation, as the average 5-year survival for breast cancers of all stages is ~ 88%.

In this manuscript we describe a linear relationship between Gibbs free energy (a measure of thermodynamic energy for a specific disease), and disability weight (a clinically validated measurement of a person’s functional status) for several neurological diseases. The choice of thermodynamics for the analysis was not fortuitous. Thermodynamic energy represents an important driving factor of chemical and biological interactions for all living organisms, and its correlation with biological events is not unexpected. Gibbs Free Energy, which incorporates information of both mRNA expression as well as protein-protein interaction, would be expected to have the ability to discriminate between “passenger genomic events” and “driving genomic events”. This remains the main quandary - finding ways to differentiate between causative vs ancillary molecular changes. The use of Gibbs Free Energy represents a novel approach, and may be of particular usefulness in neurological disorders/ neurodegerative diseases where complex and by and large unclear etiology, pathogenesis and clinical response prevent identification of a good therapeutic strategies.

Gibbs Free Energy (G) is the energy associated with a chemical reaction that can be used to do work. The free energy of a system is the sum of its enthalpy (H) plus the product of the temperature (Kelvin) and the entropy (S) of the system. We propose that the use of this well-established thermodynamic measure is useful for analyzing the interplay between patient’s genomic information and the existing knowledge about protein-protein interactions.[7] Its use is based on a number of important observations. First, proteins interacting with a large number of other proteins (even if not simultaneously) have higher entropy. Because each protein-protein interaction has a different molecular configuration, given by Boltzmann’s classic equation (S = kln(W)), the entropy, S, increases as the natural log of the number of configurations, W. As such, proteins with many interaction partners exhibit many possible configurations, and each protein-partner interaction leads to a different configuration. Highly interconnected proteins such as, for example, ubiquitin (UBC) or TP53 can undergo a simultaneous physical interaction with hundreds of their respective interaction partners, because at any given time one UBC molecule interacts with a protein and another UBC molecule interacts with another protein inside the same cell. An RNA transcriptome from a tissue biopsy thus represents an ideal mixture of UBC and its interacting partners for a given condition.

Second, transcription (RNA expression levels) data are good surrogates for protein concentration. Unlike the DNA level gene alterations, which are transcribed with variable frequencies to RNA, the number of mRNA copies is translated into individual proteins with great fidelity. Several research groups have confirmed this fidelity: Greenbaum *et al*[10] and Maier *et al*[11] report a Pearson correlation in the range of 0.4-0.9 for a large set of experiments across five different species. Similarly, Kim *et al*[12] and Wilhlem *et al*[13] found an 83% correlation between human transcription data and mass spectrometry proteomic data for multiple tissue types, supporting the use of human transcriptome as a surrogate for protein concentration.

Third, the use of real-world dataset contains an inherent level of noise, but as new mRNA data sets emerge from ongoing clinical trials, the accuracy of the information in protein-protein interaction databases as well as in the gene expression data sets will improve as well. For the time being, the preliminary findings are a great way to discover new avenues for future analysis, but as data integrity and quality of our conclusions improve, we should be able to use this data for reliable therapeutic decisions. In addition, the ability to combine different data sources (mRNA expression data from individual patients and existing PPINs) as introduced in this manuscript, is likely to deliver new insight into biologically complex diseases than traditional approaches.

## Materials and Methods

Because the study of chemical thermodynamics embodies chemical potential, for two molecules A and B interacting to form a new molecule, or an A-B molecular complex, the amount of A-B formed would be dictated by the amount of A and B. In cases where A is present in higher concentration than B, a chemical potential develops. As such, a protein, D, interacting with proteins, C, E, and F has chemical potential represented by:

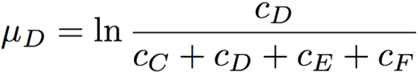

where the chemical potential is the natural log of the concentration of protein D divided by the sum of the concentrations of protein D and all the concentrations of its neighbors. Because the argument of the natural log is a ratio, we can use scaled “concentrations,” or in this case, scaled expression values. The log-2 normalized expression data typically fall in the range [−10, 10]. Rescaling sets the range to be [0,1], and we can compute the minimum e_min_, and the maximum, e_max_, for a given expression data set. The normalized expression value for each gene are then computed as follows:

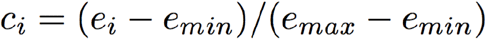

The rescaling is justified from a mathematical perspective by the fact that the argument to the natural logarithm must be positive. Furthermore, if a gene mutation leads to loss of its RNA transcription (RNA transcription is said to be down-regulated), the concentration for the respective protein would essentially be zero. Likewise, when a gene alteration leads to constitutive activation of its transcription, multiple copies of its mRNA will be made (the RNA transcription is said to be highly up-regulated), and very large quantities of protein will be produced. In this case, the protein concentration would be effectively set to the maximum of 1.

Thus, the computation of Gibbs free energy for a single protein in the PPI would be,

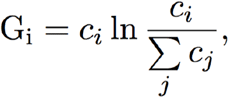

which tells us to first compute the chemical potential, for protein *i* with neighbors *j,* and multiply it by the [0,1] scaled concentration to get the Gibbs free energy for protein *i*. For the overall Gibbs Free Energy for the network, the individual G_*i*_s for each of the protein within the network are summed up. In the final calculation the normalized expression data are overlaid on the BioGRID PPI and G_*i*_ followed by use of the above equation.

We used three main data sources for our analysis: World Health Organization data on disability associated with neurological diseases, protein-protein interaction data from Biological General Repository for Interaction Datasets (BioGRID, Human ver. 3.4.139, September 2016, https://thebiogrid.org/), and RNA transcription data sets from Gene Expression Omnibus (GEO).

### WHO Tools and Statistics for neurodegenerative diseases

Unlike the terminal conditions such as cancer, there is no direct correlation between death rate and disability in neurodegenerative diseases, and death rate is not a meaningful measure of morbidity. For example, while epilepsy may cause more deaths, multiple sclerosis (MS) far outweighs its impact in the sense of morbidity and disability during a person’s lifetime. For this reason, the World Health Organization has elected Disability-Adjusted Life Years (DALYs) for evaluation of disability associated with neurodegenerative disease. DALYs combine two components: years of life lost due to premature mortality, and years lived with disability. DALYs are an expression of the number of healthy years a person looses in life. It is a more accurate representation of the damage a disease exerts on a healthy human population, and its measure - Disability Weight - is a number between 0 (perfect health) and 1 (death). While DALYs are population-dependent, and the same disease may lead to a larger apparent loss in a region where the disease is widespread, Disability Weight avoids the potential for biasing the numbers towards more prevalent diseases, and away from uncommon diseases, because it rates severity of a disease in an individual. The most recent public list of WHO for neurological diseases and their corresponding disability weights was published in 2006, and the “Neurological Disorders, public health challenges”[1] describes the demographics, geographic distributions, graphs and projections for many neurological diseases. It standardizes the comparison of neurological diseases, and was therefore used for our analysis.

### Sources of transcription data

The source of the transcription data was Gene Expression Omnibus (GEO) repository of -omics and high-throughput data https://www.ncbi.nlm.nih.gov/geo. The data sets for each of the selected neurological diseases were collapsed from probe IDs to gene IDs using GenePattern software (Broad Institute, Cambridge, MA; https://software.broadinstitute.org/cancer/software/genepattern), and the corresponding chip platform documented for the respective data set. The following datasets for specific diseases were examined: Alzheimer’s disease (GDS4136), Parkinson’s disease (GSE6613), Multiple Sclerosis (GSE19285), Epilepsy (GSE32534), Cerebrovascular disease (GSE36791), and Meningitis (GSE40586). Most data sets were already log-2 normalized, and transformed those that were not. Table 1 lists the GEO dataset number and pubmed ID (PMID) for each of the diseases.

**Table 1:** Disability score and Gibbs free energy for the neurological disorders studied

| Disease | Tissue Analyzed | GEO number | PMID | Disability Weight | Gibbs Free Energy | Number of Patients |
| --- | --- | --- | --- | --- | --- | --- |
| Alzheimer Disease | hippocampus | GSE28146 | 21756998 | 0.666 | -14735 | 22 |
| Cerebrovascular Disease | blood in ruptured intracranial aneurysms | GSE36791 | 23512133 | 0.266 | -11761 | 43 |
| Epilepsy | neocortex tissue | GSE32534 | 23418513 | 0.113 | -11467 | 5 |
| Meningitis | peripheral blood cells | GSE40586 | 23515576 | 0.615 | -16918 | 21 |
| Multiple Sclerosis | peripheral blood cells | GSE19285 | 22727118 | 0.411 | -12109 | 30 |
| Parkinson Disease | whole blood cells | GSE6613 | 17215369 | 0.351 | -10043 | 50 |

## Results and Discussion

Neurological disorders are common and represent a major public health problem. Even though neurological impairment and its sequelae constitute over 6% of the global burden of disease,[1] the management of neurological disorders has not significantly changed in the past few decades, and the mainstay of therapy remains focused on symptomatic management. As such the already very high disease burden is likely to continue to increase as the life spans across the world’s population increase. According to the recently published Global Burden of Disease 2010 Study (GBD 2010),[14] stroke is the second leading cause of death globally and the third leading cause of premature death and disability as measured in disability-adjusted life-years (DALYs).

There are many reasons for the lack of effective therapies, but the principal challenge is the complexity of data and the inconsistency in assigning causality to genomic alterations. There exists a great struggle with discriminating between incidental molecular findings, and those that may be driving disease pathogenesis. This not only hinders the search for effective therapies, but the absence of tissue targets also prevents effective clinical initiatives. Until recently much effort was dedicated to statistical interpretation of RNA expression levels. But the level of mRNA expression is not always reflective of the gene importance in a biological event on in a particular disease. An overexpressed gene that is peripheral to a major proliferative pathway will have minimal effect on proliferation, whereas a mild elevation in the expression levels of a well-connected gene will have a crucial effect on the process. Minute changes in the levels of genes coding for very important growth factors such as VEGF, or inflammation regulators interleukins lead to biological events that are normally carefully managed through feedback loops.

Any measurement of disease activity must therefore incorporate the intracellular protein-protein interactions. We introduce a method for interrogating individual tissue expression of mRNA against existing, well-curated protein-protein interaction networks. We focus on proving that thermodynamics, i.e. the molecular changes defined by Gibbs free energy, can be correlated with disease state and progression. Figure 1 shows the correlation of Disability Weight with average Gibbs Free Energy for six neurological conditions. The relationship between disease-related disability and Gibbs Free Energy is linear, with progressively worsening disability correlating with increasing more negative Gibbs Free Energy. The respective values used for the figure are shown in Table 1. We confirm that low entropy (less negative Gibbs free energy) for epilepsy correlated with the lowest disability, whereas the higher entropy (more negative Gibbs free energy) in Alzheimer’s disease correlated with the highest disability. These findings correlate with clinical observations that the severity of neurological dysfunction in multiple sclerosis, bacterial meningitis and Alzheimer disease is certainly higher than in epilepsy.

**Figure 1.**
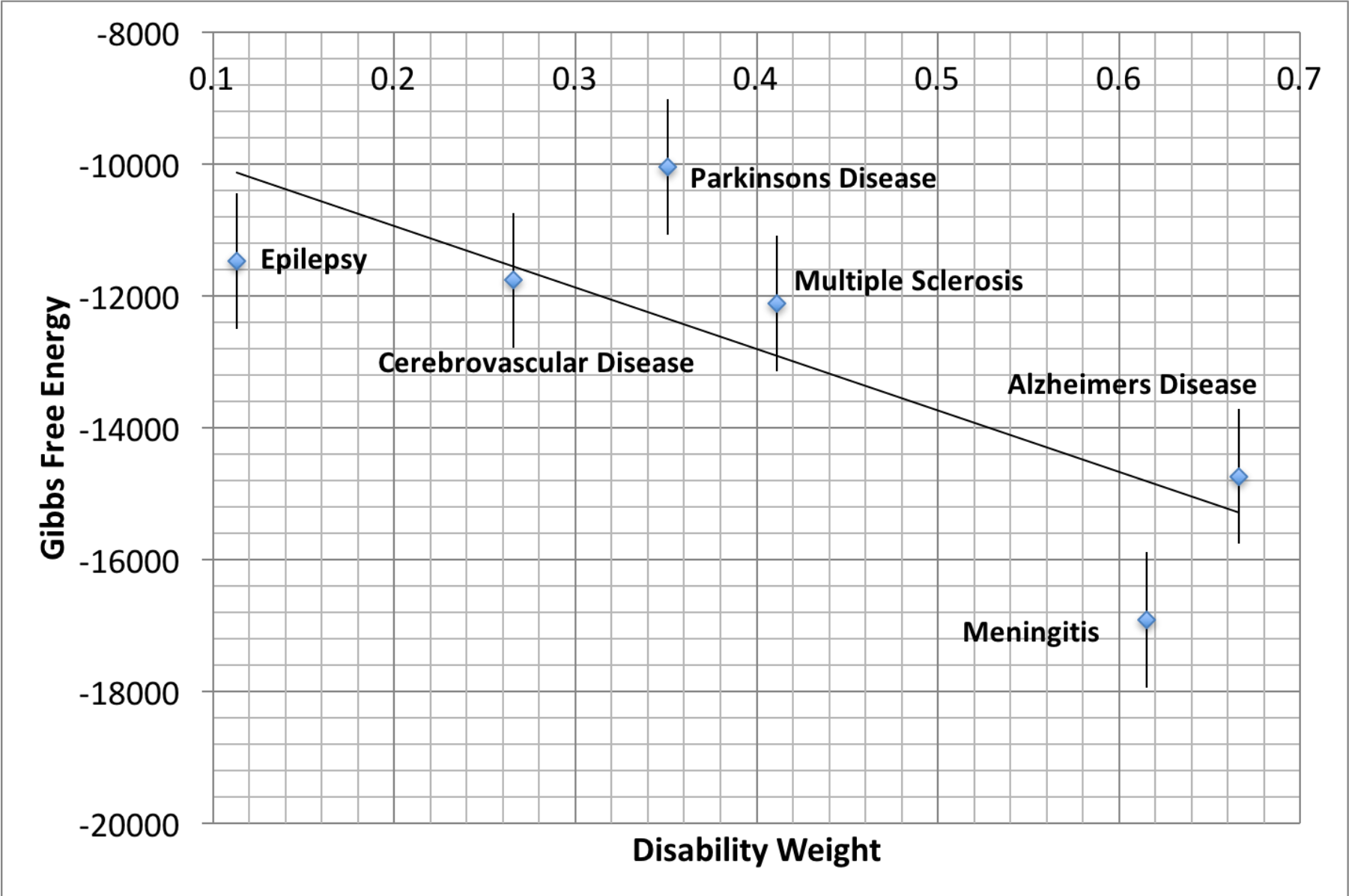
The correlation of Disability Weight and Gibbs Free Energy for six distinct neurological conditions. The mRNA expression values available from publicly available GEO data sets (Alzheimer’s disease GDS4136, Parkinson’s disease GSE6613, Multiple Sclerosis GSE19285, Epilepsy GSE32534, Cerebrovascular disease GSE36791, and Meningitis GSE40586) were used to calculate Gibbs free energy, a thermodynamic measure of protein-protein interactions. As would be expected based on the level of clinical disability, epilepsy has low Gibbs free energy (low entropy) and correlates with the lowest neurological disability. This is in stark contrast with the high Gibbs Free Energy (high entropy) and high neurologic disability in Alzheimer disease as would be consistent with clinical observations in this disease. The respective values, and the size of cohort are summarized in Table 1, and the error bars have been set to 5% of the average Gibbs Free energy value, given that the actual errors of the reported mRNA gene expression values were not reported.

We further explored whether Gibbs free energy correlated not only with severity of the disease, but also with disease progression. Seven stages have been described in the Alzheimer Disease (AD) progression, from no impairment (Stage 1) to loss of ability to respond to their environment or communicate, needing assistance with all activities of daily living, and loss of ability to swallow (Stage 7). The GEO data sets tend to simplify these stages into only 4 stages (Figure 2a), and we show a clear linear relationship (R = 0.7978) between thermodynamic energy (Gibbs free energy) and the 4 stages of the disease. The positivity of the slope is most likely reflective of neuronal loss, with less and less metabolically active tissue. This is in direct contrast with epilepsy, typified by an abnormality of conduction of an action potential across neuronal tissues rather than by a neuronal loss, where Gibbs free energy is less negative (i.e. more positive).

**Figure 2.**
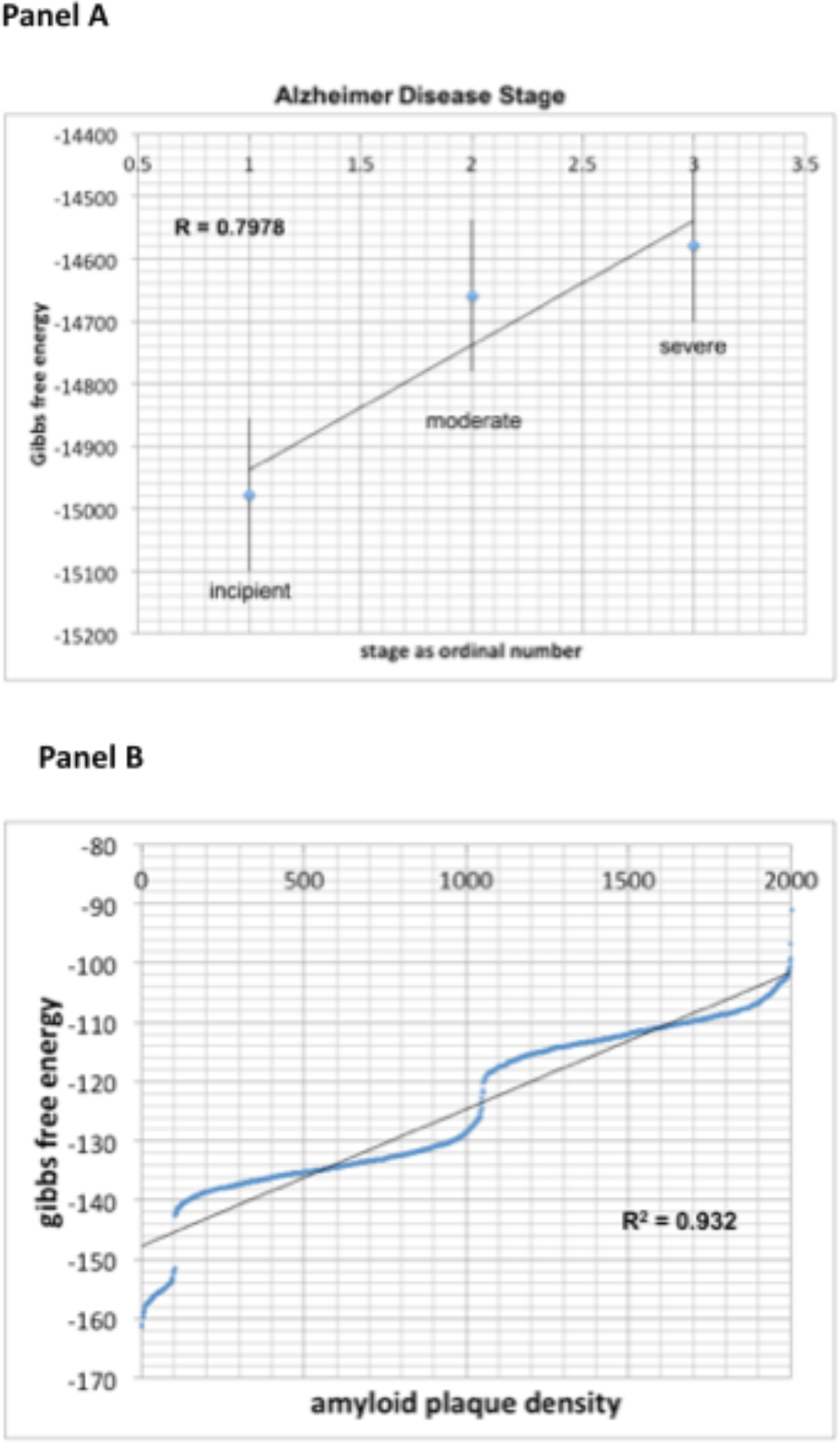
The correlation between disease progression and pathological findings with Gibbs free energy in Alzheimer Disease. The analysis of 2000+ Alzheimer Disease patients from GSE84422 data set revealed a correlation coefficient R = 0.7978 between Gibbs Free Energy and disease progression using disease stage (panel a), and an even stronger correlation (R= 0.932) using tissue pathological analysis such as amyloid plaque density (panel b).

Similarly strong correlation (R=0.932) was observed between Gibbs free energy and the degree of amyloid deposition in Alzheimer disease (Figure 2b). In this case, progressive histological changes in the form of extracellular deposits of amyloid β peptides, senile plaques, and intracellular neurofibrillary tangles of hyperphosphorylated tau in the brain, relate to neuronal death and correlate with severe disability. The amyloid metabolic cascade and the posttranslational modification of tau protein, often considered causal in AD, are sufficient to explain the diversity of biochemical and pathological abnormalities in AD. There is a multitude of cellular and biochemical changes leading to the accumulation of extracellular senile plaques made of deposits of Aβ peptide, and all have the outcome of neural degeneration and loss. The correlation of Gibbs free energy with histological changes and disease progression implies a global value of our measurement and the need for future evaluation of the hidden metabolic information.

Yet another scale is used by clinicians for evaluation of disability in multiple sclerosis (MS). The Expanded Disability Status Scale (EDSS), was developed by a neurologist (John Kurtzke) in 1983, and ranges between 0-10. It is based on neurological evaluation of pyramidal (limb movement), cerebellar (ataxia, coordination, tremor), brainstem (speech, swallowing and nystagmus), sensory, bowel and bladder, vision, or cerebral (mental) functions. Its 0.5 unit increments represent progressively higher levels of disability. EDSS 1.0-4.5 refers to people with MS who are able to walk without any aid, whereas 5-6 refers to progressive motor and cognitive disability. Similarly to AD, we find a positive linear slope between Gibbs Free Energy and disability in MS (Figure 3), as consistent with neuronal loss due to demyelination. The respective correlation coefficient between EDSS and Gibbs free energy in MS was found to be very strong at R=0.913.

**Figure 3.**
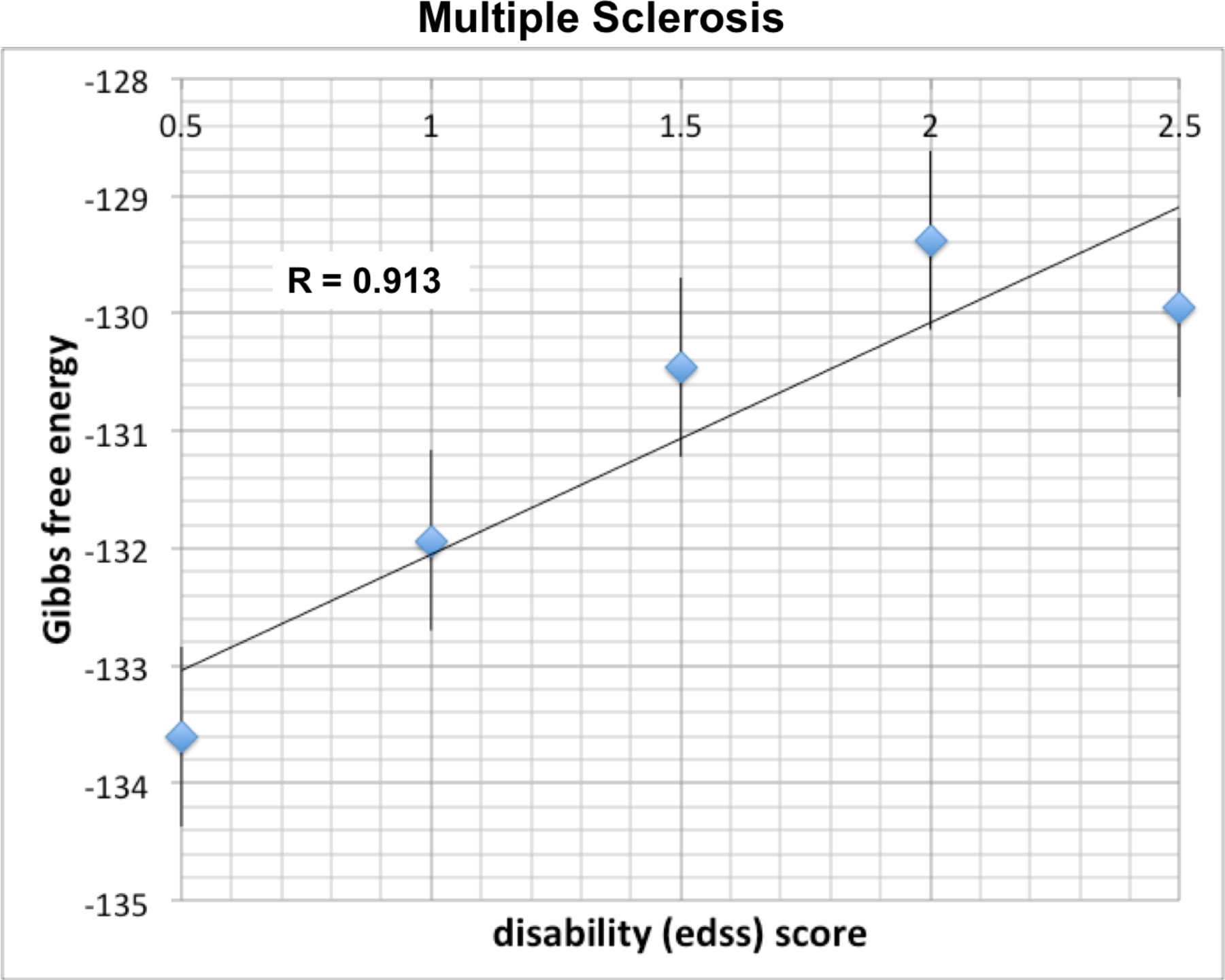
The correlation between disease progression and Gibbs free energy in Multiple Sclerosis. Thirty patients with multiple sclerosis from the GEO data set GSE19285 revealed a correlation coefficient R=0.913 between Gibbs Free Energy and disease progression as reflected by The Expanded Disability Status Scale (EDSS).

## Conclusions

We provide early evidence for using not only expression data, but also connectivity/topology data for analysis of genomic information in neurodegenerative disease. One of the approaches is using a thermodynamic measure such as Gibbs Free Energy to identify the disease-related global changes in protein-protein interactions network resulting from changes in genomic profiles of individual patients. The ability to correlate these molecular profiles disease severity and disease progression suggests that we can use Gibbs Free Energy in the future to evaluate causative vs ancillary molecular changes through mathematical simulation of the protein inhibition/stimulation. This is the first installment in developing mathematical algorithms that would facilitate identification of relevant, therapeutically targetable pathways in neurodegenerative diseases. The use of Gibbs Free Energy in genomic analysis would be beneficial not only from therapeutic point of view, but also from cost and sustainability perspective, because it would minimize futile clinical trials.

## Acknowledgments

We acknowledge (NSF)/ECCS-1533693 NSC-FO: Col “Individual Variability in Human Brain Connectivity, Modeling Using Multi-scale Dynamics Under Energy Constraints”, and Office of Naval Research (ONR)/N00014-15-2126. The content is solely the responsibility of the authors and does not necessarily represent the official views of the National Science Foundation, or the US Navy. We thank Samuel McGuire for helpful discussions. EAR and GLK acknowledges partial funding support from CSTS Healthcare, Toronto, Canada.

